# Suppression weakens unwanted memories via a sustained reduction of neural reactivation

**DOI:** 10.1101/2021.01.16.426815

**Authors:** Ann-Kristin Meyer, Roland G. Benoit

## Abstract

Aversive events sometimes turn into intrusive memories. However, prior evidence indicates that such memories can be controlled via a mechanism of retrieval suppression. Here, we test the hypothesis that suppression exerts a sustained influence on memories by deteriorating their neural representations. This deterioration, in turn, would hinder their subsequent reactivation and thus impoverish the vividness with which they can be recalled. In an fMRI study, participants repeatedly suppressed memories of aversive scenes. As predicted, this process rendered the memories less vivid. Using a pattern classifier, we observed that suppression diminished the neural reactivation of scene information both globally across the brain and locally in the parahippocampal cortices. Moreover, the decline in vividness was associated with reduced reinstatement of unique memory representations in right parahippocampal cortex. These results support the hypothesis that suppression weakens memories by causing a sustained reduction in the potential to reactivate their neural representations.

## Introduction

Memories of the past are not always welcome. There are experiences that we would rather not think about, yet that involuntarily intrude into our awareness. Research over the last two decades has demonstrated that we are not at the mercy of such unwanted memories: we can control them by actively suppressing their retrieval (1–3). This process weakens the memory and can eventually cause forgetting (4). Here, we seek to tie the sustained phenomenological weakening of a suppressed memory to its neural basis.

Neuroimaging research has made strides in determining the transient neural mechanisms that prevent unwanted retrieval. It has consistently shown that retrieval suppression is associated with increased activity in right dorsolateral prefrontal cortex (dlPFC) and decreased activity in the hippocampus (5–13). This pattern has been interpreted as a top-down inhibition of critical hippocampal retrieval processes by the dlPFC (5–7).

During retrieval, the hippocampus is integral for reinstating the cortical activity patterns that were present during the encoding of the memory (14–18). Inhibition of the hippocampus would accordingly hinder such momentary cortical reactivation and thus prevent unwanted retrieval (10, 19). Consistent with this account, retrieval suppression has been found to also affect activity in cortical regions that encode the particular content of the suppressed memory (e.g. 6, 9–11, 13).

For example, when the unwanted memories comprise images of complex scenes, suppression is accompanied by a transient reduction of activation in the parahippocampal cortex (PhC) (8). This region particularly supports memories for scenes (20–23) and its activity during retrieval scales with the detailedness (24, 25) and vividness (26–28) of the memories. Moreover, more fine-grained analyses of the activity patterns within the PhC have linked the reactivation of memory-specific representations to the successful retrieval of scenes (21, 29, 30). In turn, there is also some evidence that attempts to suppress an unwanted memory indeed momentarily prevent such memory-specific reactivation (10, 31–33).

We have thus gained an evolved understanding of the mechanisms that are engaged transiently *during* the suppression of unwanted memories. By contrast, there is little evidence for the sustained neural *after-effect* of this process: Why do previously suppressed memories remain difficult to recall? Suppression has been argued to deteriorate the memory’s neural representation (19, 34). Here, we test the hypothesis that it thus compromises later reactivation, even when one then tries to intentionally recall that memory (see also 35). A deficient reactivation of PhC representations would hinder such recall attempts and diminish the vividness of the recollection.

To test this hypothesis, we conducted an fMRI study using an adapted *Think/No-Think* procedure (36, 37). First, participants learned associations between neutral objects (cues) and aversive scenes (target memories) (Fig. 1a). During the suppression phase, they were then scanned by fMRI while they again encountered the cues. In this phase, participants were repeatedly prompted to recall the associated target for one third of the cues (*recall condition*), whereas they were requested to prevent the retrieval of the targets for another third of the cues (*suppress condition*). In the suppress condition, we instructed participants to remain focused on the cue while trying to block out all thoughts of the accompanying target memory without engaging in any distracting activity (7, 38). Importantly, the remaining third of the cues were not presented during this phase (*baseline condition*). These cues and their associated targets thus serve as a baseline for the passive fading of memories that simply occurs due to passage of time (i.e., without any active suppression attempts).

**Fig. 1.**
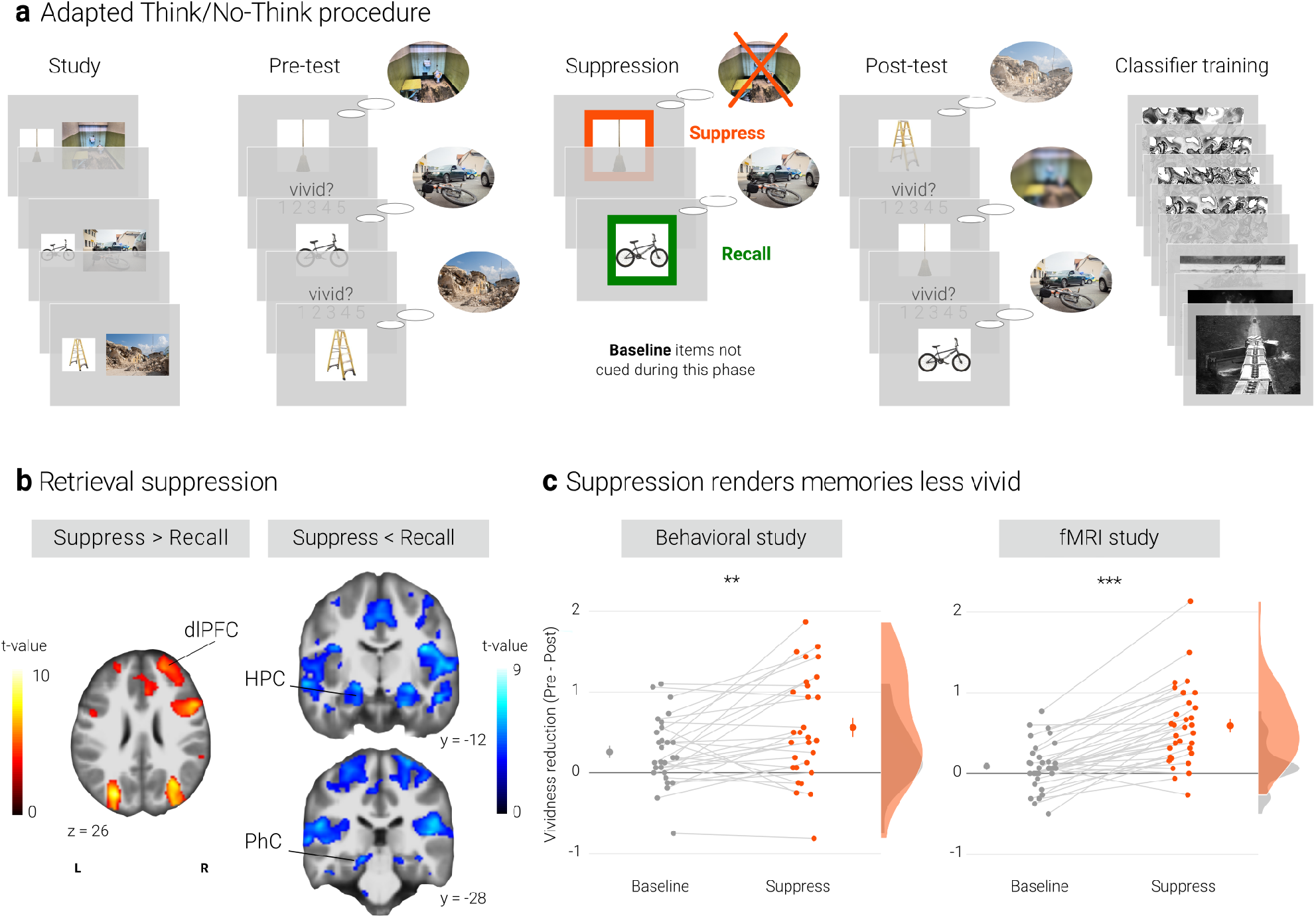
Experimental procedure, univariate MRI and behavioral results. **a)** Illustration of the adapted Think/No-Think procedure. Participants studied associations between unique objects and aversive scenes. Both during a pre- and a post-test, they covertly recalled all the scenes in response to the objects and rated the vividness of their recollection. In between these two tests, they performed the suppression phase. Specifically, for objects presented in a green frame, participants repeatedly recalled the associated scene (*recall condition*). By contrast, for objects presented in a red frame, they suppressed the retrieval of the associated scene (*suppress condition*). Note that we did not present a third of the objects during this phase (*baseline condition*). Finally, participants performed a one-back task that served to train a pattern classifier in detecting evidence for scene reactivation. **NB** Following the IAPS user agreement, we have replaced the original pictures with similar scenes for this figure. As per bioRxiv policy, we further replaced scenes featuring people by stand-ins. In the original stimulus set, each object cue also features in its paired scene. **b)** The suppression phase yielded the typical activity pattern associated with retrieval suppression, including greater activation in the right dorsolateral prefrontal cortex (dlPFC) and reduced activation in the hippocampus (HPC) and parahippocampal cortex (PhC) during *suppress* versus *recall* trials. For display purposes, the images are thresholded at *p* < .001, uncorrected, with a minimum cluster size of 50 voxels. **c)** Suppression caused a reduction in self-reported vividness from the pre-to post-test that exceeded any change due to the passage of time as indexed by the *baseline* condition. This effect replicated across the fMRI study (*n* = 33) and a behavioral study (*n* = 30) with an independent sample. Large dots indicate the mean, error bars the standard error of the mean. *** *p* < .001, ** *p* < .01.

To assess the degradation of neural memory representations over time, we also had participants recall each target in response to its cue both before and after the suppression phase. During these pre- and post-tests, they reported the vividness of the recalled memories. We thus assessed the phenomenological quality of the memories at the same time that we probed their neural reinstatement. Finally, participants engaged in a separate task that allowed us to train a pattern classifier to detect neural reactivation of complex and aversive scenes.

We tested our hypothesis by tracking the impact of suppression on neural reinstatement. Specifically, we tested four key predictions. First, we expected that suppression would be associated with reduced scene reactivation, both distributed across the brain and more regionally specific in the PhC (see also 10, 31, 33). Second, we predicted that this effect would not be confined to the transient moment of active suppression but also linger on - as indexed by lower post-test reactivation of previously suppressed memories. Third, in addition to a reduced reactivation of general scene information, we also predicted a weaker PhC reinstatement of the neural representations that are unique to the individual memories. Finally, if weaker neural reactivation constitutes the basis for the sustained suppression-induced reductions in vividness, we expected a relationship between these effects.

## Results

### Preventing retrieval yields the typical pattern associated with memory suppression

We first sought to establish whether our procedure elicited the activation pattern that has consistently been associated with retrieval suppression (e.g. 7, 13, 19, 39). Suppressing versus recalling an aversive scene indeed led to increased activation in a number of brain regions including the right dlPFC and reduced activation in, amongst others, the bilateral hippocampi and PhC (Fig. 1b, Supplementary material). This pattern is consistent with the engagement of the mechanism thought to mediate retrieval suppression (7, 10).

In the following, we test the hypothesis that this mechanism impairs subsequent retrieval attempts by hindering reinstatement of the neural memory representation. We thus examine suppression-induced changes in the phenomenological quality of the memories and their neural basis. These analyses focus on the critical comparison of the *suppress* versus *baseline* conditions. In the Supplementary material, we explore possible effects of retrieval practice (40–42), i.e., contrasts of the *baseline* versus *recall* conditions.

### Suppression renders memories less vivid

We assessed the impact of suppression on the phenomenological quality of the memories by examining their change in vividness from the pre-test to the post-test. Indeed, there was a greater reduction for *suppress* than *baseline* memories as indicated by a significant interaction between time of test (pre, post) and condition (*baseline*, *suppress*) (*F*(1, 32) = 46.18, *p* < .001, 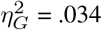). However, the main effects of time of test (*F*(1,32) = 28.87, *p* < .001, 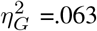) and condition (*F*(1,32) = 4.22, *p* = .048, 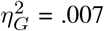) were also significant. Follow-up tests showed that suppression reduced the vividness of the memories (*t*(32) = 6.60, *p* < .0001, *d* = 1.17), whereas baseline memories did not significantly change over time (*t*(32) = 1.79, *p* = .08,, *d* = 0.32) (Fig. 1c).

We obtained largely the same pattern in a behavioral study with an independent sample (Fig. 1c): again, the critical interaction between time of test and condition was significant (*F*(1, 28) = 8.85, *p* = .006, 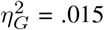). Additionally, there were main effects of time of test (*F*(1, 28) = 21.78, *p* < .001, 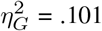) and of condition (*F*(1, 28) = 8.02, *p* = .008, 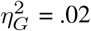). Baseline and suppress memories did not differ on the pre-test (*t*(28) = 0.48, *p* = .63, *d* = 0.09) but on the post-test (*t*(28) = 3.26, *p* = .003, *d*= 0.62). However, the follow-up tests showed a reduction in vividness for suppressed (*t*(28) = 4.63, *p* < .0001, *d* = 0.88) and also for baseline memories (*t*(28) = 3.41, *p* = .002, *d* = 0.64).

Consistent with prior research (4), suppression thus had a replicable, detrimental impact on people’s ability to vividly recall the suppressed memories. Importantly, we assessed the phenomenological quality of the memories during exactly those retrieval attempts that also provide the basis for our critical fMRI analyses. That is, in the following, we examine not only whether there is less reactivation of a memory during suppression (10, 31), but also the hypothesis that this effect then lingers on during these subsequent recall attempts.

### Establishing a linear classifier to detect scene reactivation

Memory retrieval reactivates the perceptual and conceptual representations elicited during encoding (43, 44). To quantify the degree of such reactivation on a given trial, we trained a linear support vector machine (45) on data from an independent task that participants had performed at the end of the MRI session. Specifically, the classifier learned to distinguish brain states associated with the perception of intact aversive scenes (similar to the ones used in the main task) versus morphed versions of the scenes. The morphed scenes were created via a diffeomorphic transformation that renders them unrecognizable while preserving their basic perceptual properties. Compared to conventional methods, such as scrambling, morphing has been shown to elicit neural activation that is more similar to activation induced by intact images (46).

Given the widespread nature of memory representations (17, 47, 48), we sought to test for global reactivation by training a classifier on all voxels of the respective participant’s grey matter mask. Using cross-validation, the classifier reached a mean accuracy of 80.3% (*SD* = 17.4) on the training data, corroborating that it was able to distinguish brain states associated with the presentation of intact versus morphed aversive scenes (*t*(32) =10, *p* < .001). We then used the trained classifier to analyze the activity patterns on each trial of our memory tasks. Specifically, for a given trial, we calculated the dot product of the trial’s activation map and the classifier’s weight pattern (49, 50). We take the resulting values to index the degree of scene reactivation (Fig. 2a). (To ensure that the effects obtained with this mask are not simply driven by the PhC, we also ran all analyses for an additional ROI that excluded this region from the whole-brain mask. The results were virtually identical to the ones reported throughout the manuscript as described in the R Markdown available at OSF (https://osf.io/27dkh/?view_only=8ec25028113941f683f31703ab580533).

**Fig. 2.**
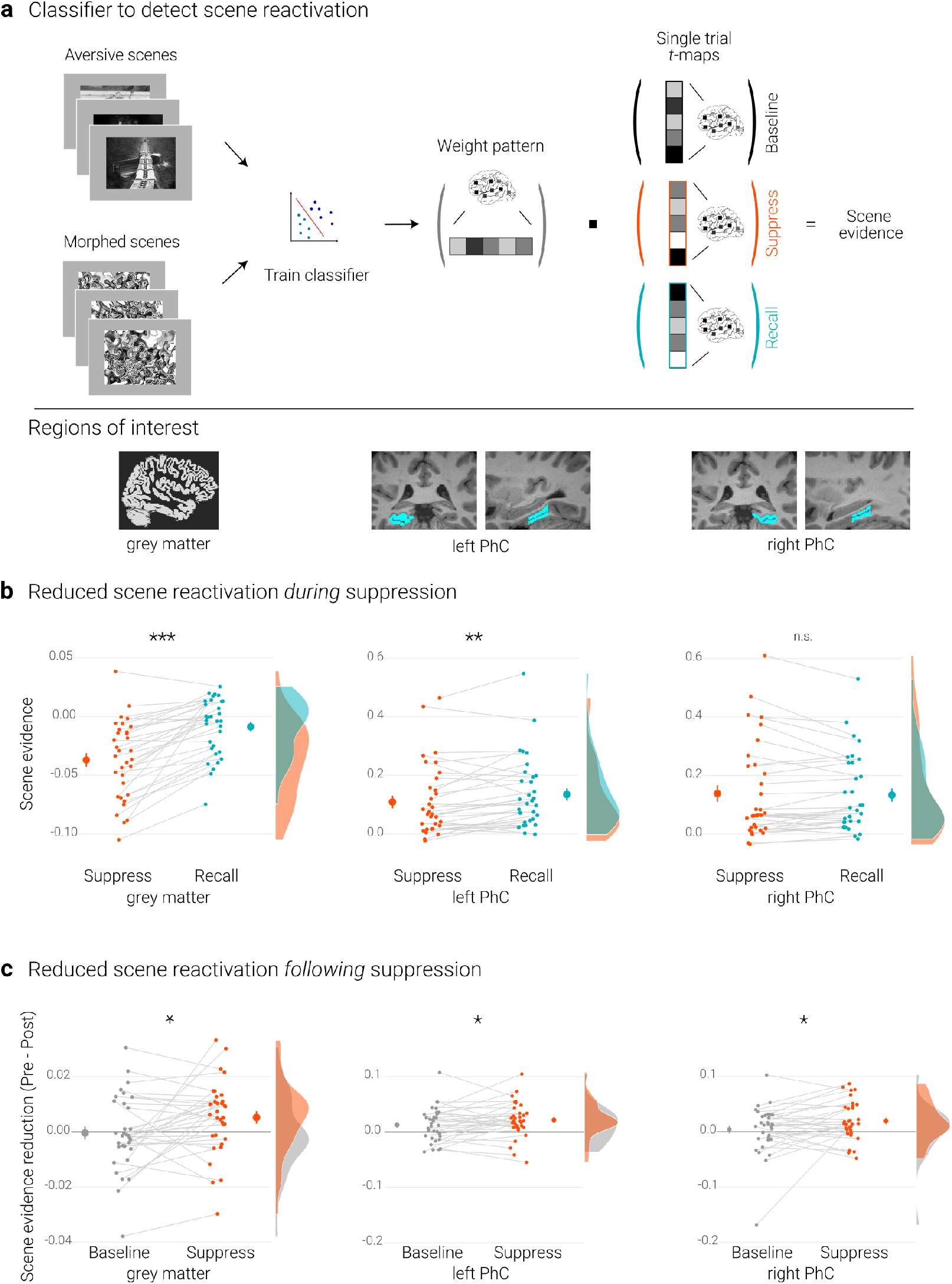
Effects of suppression on scene reactivation. **a)** A linear support vector machine was trained on data of an independent task to discriminate neural activity patterns associated with the perception of intact versus morphed aversive scenes. **NB** Following the IAPS user agreement, we have replaced the original pictures with similar scenes for this figure. As per bioRxiv policy, we further replaced scenes featuring people by stand-ins. The dot product of the resulting weight pattern and single-trial *t*-maps was used as a proxy for reactivated scene information. We compared such scene evidence between conditions globally across the grey matter and more locally in the left and right parahippocampal cortices (PhC, manually segmented on the individual structural images). **b)** Across the grey matter and locally in left PhC, there is less scene evidence while participants suppress than recall scene memories. **c)** The suppression-induced reduction in scene evidence lingers on after suppression: scene evidence decreases from the pre-to the post-test for suppressed memories but not for baseline memories. This was the case across the grey matter and in the PhC. Larger dots indicate the mean, error bars the standard error of the mean. *** *p* < .001, ** *p* < .01, * *p* < .05, *n* = 33.

We also sought to test for more localized reactivation of scene information in the PhC, given the preferential engagement of this region for scene memories (20, 51). Towards this end, we manually traced the parahippocampal cortices on each individual anatomical scan (20, 52, 53) (Fig. 2) and trained classifiers separately for the masks from the left and right hemisphere. These classifiers reached average cross-validation accuracies of 77.3% (*SD* = 16.6; *t*(32) = 9.5, *p* < .001) and 82.6% (*SD* = 16.6; *t*(32) = 11.3, *p* < .001), respectively.

We further validated our approach by examining the correspondence between the reactivation scores in the PhC and the vividness with which the memories could be recalled. The analysis was conducted on data from the pre-test. Because memories at that stage are still unconfounded by possible effects of the subsequent experimental manipulation, this allowed us to compute correlations based on all trials across the three conditions. Specifically, we correlated the scene reactivation and vividness scores for each participant and then performed one-sample *t*-tests on the individual Fisher-transformed correlation coefficients. These analyses showed that greater scene reactivation was indeed associated with more vivid recollections in left (*M* = 0.09, *95% CI* = [0.03, 0.16], *t*(32) = 2.82; *p* = .01) and right PhC (*M* = 0.06, *95% CI* = [0.01, 0.12], *t*(32) = 2.27; *p* = .03).

### Reduced scene reactivation *during* suppression

The previous section established that the classifier provides a measure for the reactivation of scene information. We first examined whether such reactivation is reduced while participants intentionally try to suppress rather than to recall a memory. This was the case globally across the brain as indicated by the analysis based on the grey matter mask (*t*(32) = 7.04, *p* < .001, *d* =1.22).

For the PhC, a rANOVA with the factors hemisphere (left, right) and condition (recall, suppress) revealed an interaction of hemisphere and condition (*F*(1,32) = 30.04, *p* <.001, 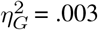). Follow-up tests showed reduced scene evidence locally in the left (*t*(32) = 2.84, *p* = .01, *d* = 0.50), though not right PhC (*t*(32) = −0.60, *p* = .56, *d* = −0.11) (Fig. 2b). These data suggest that participants were successful at controlling the retrieval of unwanted memories. At the same time, they further validate the use of the classifier as a measure of memory reactivation.

### Reduced global scene reactivation *following* suppression

Suppressed scenes were recalled less vividly than baseline scenes. We had hypothesized that this suppression-induced decline of the memories reflects a sustained reduction in the potential to reactivate their neural representations. We thus expected reactivation scores for *suppress* memories to decline from the pre-test to the post-test to a larger degree than for *baseline* memories.

We tested for this effect by conducting a rANOVA on the global reactivation scores with the factors time of test (pre, post) and condition (suppress, baseline). This analysis yielded the expected significant interaction (*F*(1,32) = 5.14, *p* = .03, 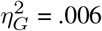), reflecting diminished scene reactivation for suppressed (*t*(32) = 2.26, *p* = .03, *d* = 0.4) but not for baseline memories (*t*(32) = −0.2, *p*= .84, *d* = −0.03) (Fig. 2c).

### Reduced parahippocampal scene reactivation *following* suppression

As predicted, suppression also lead to a sustained reduction of local scene reactivation in the PhC. This was corroborated by a rANOVA with the factors time of test (pre, post), condition (baseline, suppress), and hemisphere (left, right) that yielded the significant interaction between time and condition (*F*(1,32) = 4.33, *p* = .046, 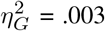) (in addition to a main effect of time, *F*(1,32) = 8.83, *p* = .006, 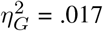). This effect reflected the expected reduction in scene reactivation for suppressed (*t*(32) = 3.77, *p*< .001, *d* = 0.67) but not for baseline memories (*t*(32) = 1.38, *p*=.18, *d* = 0.24) (Fig. 2c).

### A link between suppression-induced reductions in scene reactivation and vividness

Activity in the PhC has previously been associated with the number of details (24, 25) and the vividness (26–28) with which scenes can be recalled. We similarly observed that the recall of more vivid memories is accompanied by greater evidence for scene reactivation.

We accordingly hypothesized that a greater suppression-induced reduction in scene reactivation would lead to a greater reduction in vividness. We examined this hypothesis by exploiting the natural variation in people’s ability to control unwanted memories.

For each participant, we quantified the suppression-induced reductions in vividness as the change from the pre-to the post-test for suppressed memories, corrected for by the change in vividness for baseline memories:

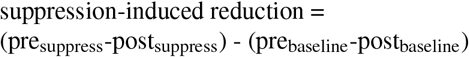

We thus obtained an index of the deterioration in vividness that exceeds any effects that simply occur due to the passage of time (7, 19). Analogously, we calculated the degree of suppression-induced reductions in scene reactivation by subtracting the change score of the baseline memories from the score of the suppressed memories.

If the reduction in reactivation is linked to the reduction in vividness, we expected a positive correlation between the behavioral and neural suppression-induced reduction scores. Indeed, using robust skipped Spearman’s correlations, we found a significant effect for the right (*r* = .46, *95%-CI* = [.08 .76]) and a trend for the left PhC (*r* = .34, *95%-CI* = [-.05 .67], Fig. 3).

**Fig. 3.**
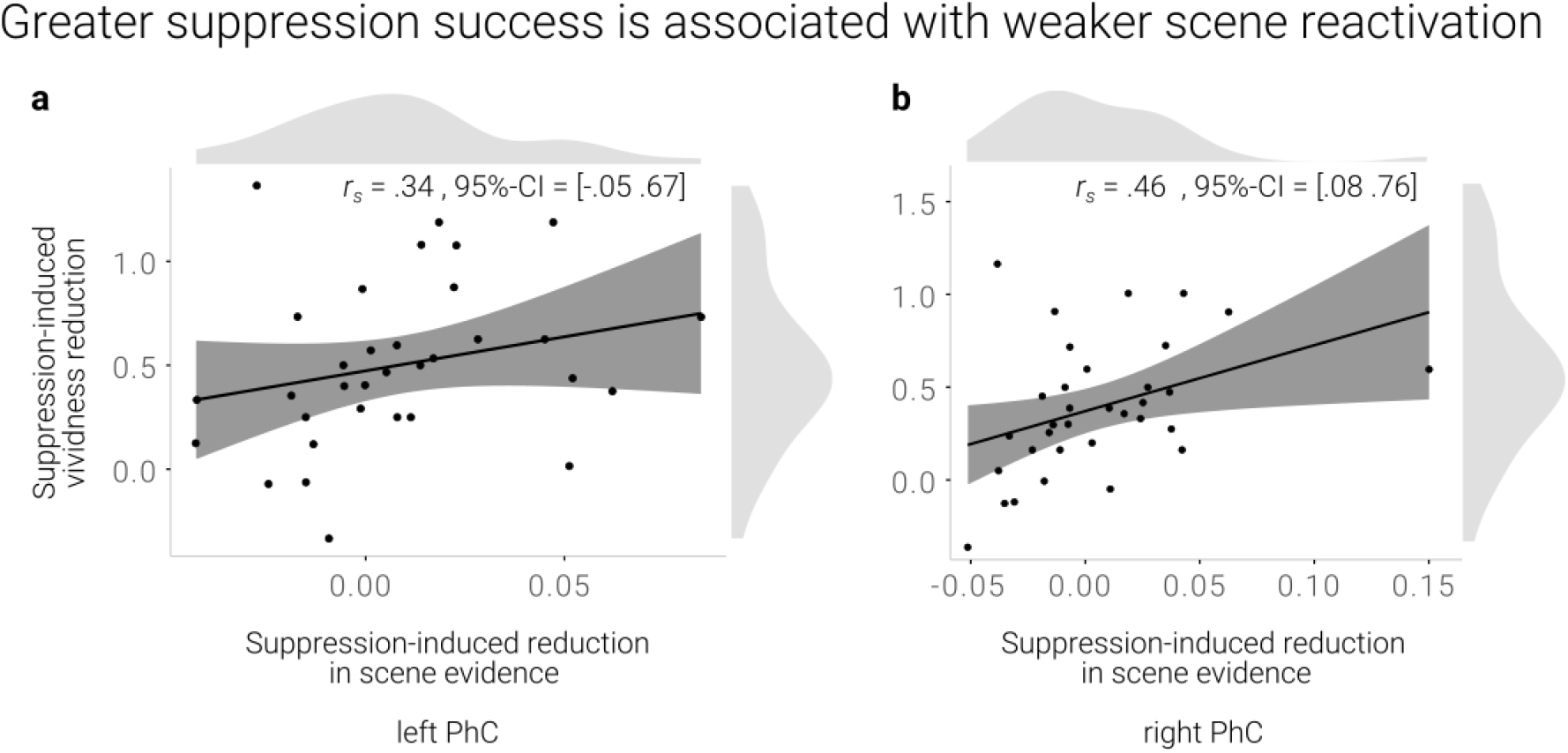
**a,b)** A greater suppression-induced reduction in vividness is associated with a greater suppression-induced reduction in scene evidence in right PhC as indicated by a robust skipped Spearman’s correlation. The left PhC showed a non-significant trend only for this effect. Black lines indicate linear regression lines, dark grey shades indicate 95% - confidence intervals, PhC: parahippocampal cortex.

Taken together, suppression led to a sustained reduction of scene information on a global and local level. Moreover, the degree of reduced reactivation of scene information in the right PhC was linked to the decline of the memories’ vividness. These data may suggest that reduced reactivation reflects the failure to retrieve scene features that would have made the recollections more vivid. In the following, we further examine this interpretation by assessing changes in the neural reinstatement of individual memory representations.

### Suppression success is associated with weaker memory-specific PhC pattern reinstatement

The classifier results indicate that suppression hinders the subsequent reactivation of scene information. However, they do not address the question whether this effect reflects reduced reinstatement of information that is specific to a particular memory. In a next step, we thus used Representational Similarity Analysis (RSA) (54, 55) to examine the reinstatement of activity patterns that are unique to the individual memories. We focus this analysis on the PhC, where the neural reinstatement of a particular memory should yield a unique and replicable activity pattern (21, 29, 30). Specifically, we expected a similar activity pattern to emerge whenever participants recall the same scene memory.

We quantified similarity by computing the Pearson correlation (54, 55) of the activity patterns across the pre- and post-test. As an index of memory-specific reinstatement, we then compared the similarity of a memory with itself (*same-item similarity*) and the similarity of a memory with all other memories of the same condition (e.g., baseline) (*different-item similarity*) (29, 56, 57) (Fig. 4a).

**Fig. 4.**
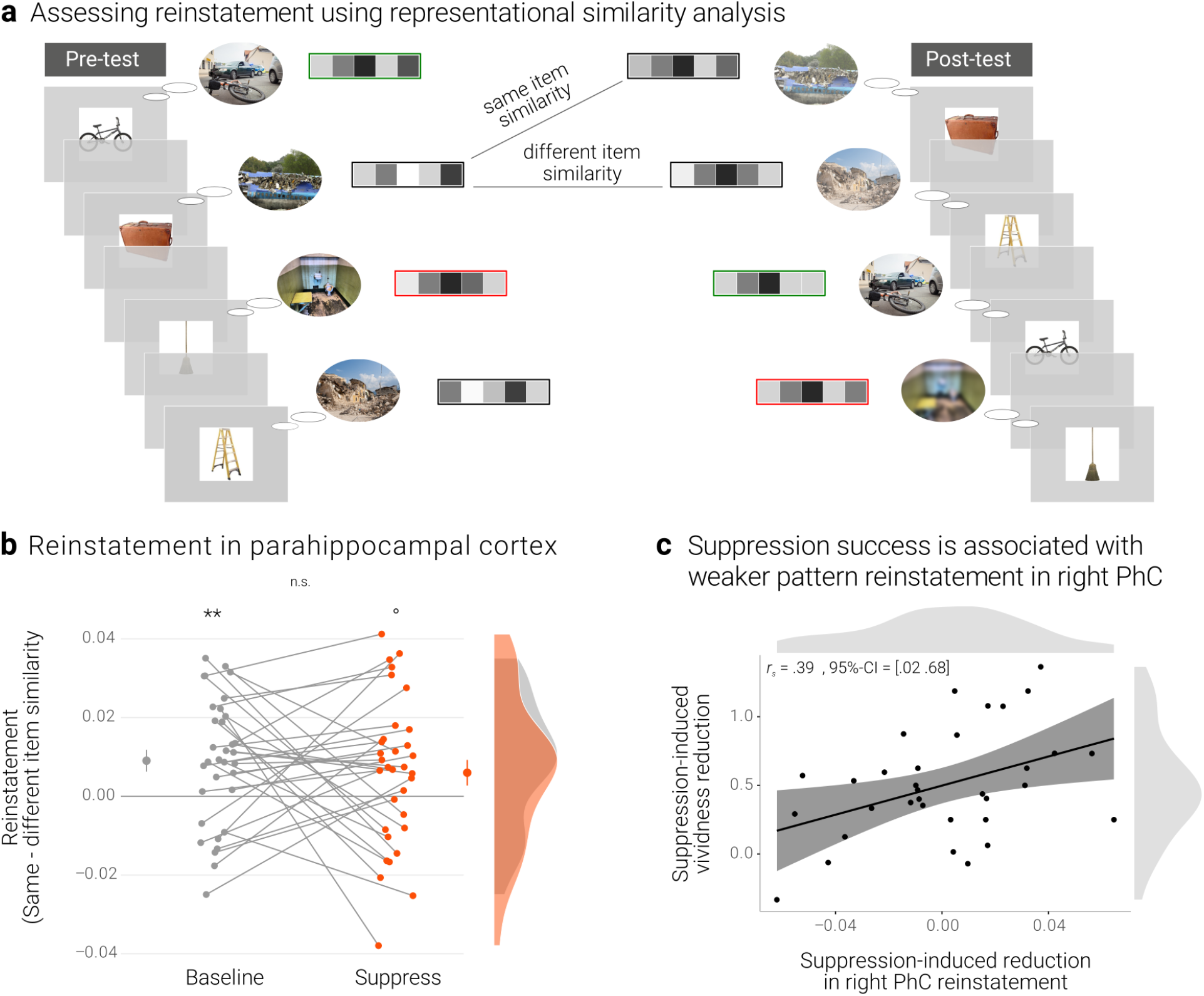
Effects of suppression on individual memory representations. **a)** We estimated the reinstatement of the neural memory representations by assessing the similarity of the activity patterns across the pre- and the post-test. We take the difference between the same-item and different-item similarity as an index of neural reinstatement. **NB** Following the IAPS user agreement, we have replaced the original pictures with similar scenes for this figure. As per bioRxiv policy, we further replaced scenes featuring people by stand-ins. **b)** In the PhC, the difference between same- and different-item similarity was significant for the baseline memories. These memories thus seem to have been consistently reinstated across the two tests. By contrast, the suppressed memories only showed a trend for this effect, though the critical interaction was not significant. Large dots indicate the mean, error bars the standard error of the mean. **c)** A greater suppression-induced reduction in vividness was associated with a greater suppression-induced reduction in reinstatement in the right PhC (as indicated by a robust skipped Spearman’s correlation). Black lines indicate linear regression lines, dark grey shades indicate 95% - confidence intervals, PhC: parahippocampal cortex. ** *p* < .01, * *p* < .05, ° *p* < .1, *n* = 33.

However, we note that the scene memories were probed with the same objects on both the pre- and post-test. Though the PhC is more sensitive to scene than object information (20), any difference in same-versus different-item similarity may thus partly reflect the repetition of these retrieval cues. Critically, this caveat would not explain any differences in reinstatement for baseline versus suppressed memories.

A rANOVA with the factors scene identity (same, different), condition (baseline, suppress), and hemisphere (left, right) yielded the main effects of identity (*F*(1,32) = 13.64, *p* < .001, 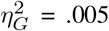) and hemisphere (*F*(1,32) = 5.41, *p* = .027, 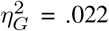) though not the critical interaction between scene identity and condition (*F*(1,32) = 0.46, *p* = .501, 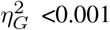). In follow-up analyses, we observed greater same-than different-item similarity for the baseline memories (*t*(32) = 3.25, *p* = .003, *d* = 0.58) but only a nonsignificant trend for a numerically smaller effect for the suppress memories (*t*(32) = 1.84, *p* = .075, *d* = 0.32) (Fig. 4b). The data thus provide some evidence for the replicable reinstatement of neural representations that are unique to the individual memories. Though there was only a trend for this effect in suppressed memories, it was - overall - not significantly smaller than for the baseline memories.

We had hypothesized that individuals who were more successful at suppression (as indicated by a greater reduction in vividness) should show evidence for a greater decline in neural reinstatement. As for the reactivation scores above, we thus examined the association between the behavioral suppression-induced reduction scores and the difference in reinstatement for baseline versus suppressed memories. The latter was computed as

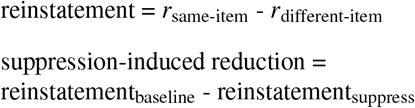

Thus, a greater value indicates a greater reduction in memory-specific reinstatement. Mirroring the results of the pattern classifier, the skipped Spearman’s correlation between the behavioral and neural effects was significant in the right (*r* = .39, *95%-CI*= [.02 .68]) though not left PhC (*r* = .02, *95%-CI* = [-.37 .41]) (Fig. 4c).

## Discussion

Research over the last two decades has demonstrated that we are able to control our unwanted memories by intentionally suppressing their retrieval (1, 36). This process weakens the avoided memory and can eventually lead to forgetting (4, 37). Though this research has made strides in elucidating the transient mechanisms engaged *during* retrieval suppression (5–13), there is little evidence for the neural consequences that underlie the sustained subsequent changes in the retrievability of a suppressed memory.

In this study, we sought to tie the sustained phenomenological weakening of a suppressed memory to its neural basis. Successful episodic memory retrieval entails the reinstatement of a memory’s representation (16, 18, 43, 58, 59). It is fostered by hippocampal processes that complete the neural pattern of the original experience (e.g., of a particular scene) from a partial pattern provided by an adequate retrieval cue (e.g., of an object that was also part of the scene) (14, 15, 60).

This process leads to the cortical reinstatement of a memory across the regions that had been involved in its original encoding (23, 47, 61).

Reinstatement has been examined in humans using fMRI by exploiting the distributed pattern of activity across voxels as a proxy of a memory’s neural representation. Successful retrieval (e.g., of a particular scene) has thus been shown to be accompanied by a reactivation of categorical information (e.g., scene information) that is both widely distributed (18, 47, 48, 62) and localized to specific brain regions (e.g., the PhC; 29, 30).

In the current study, the degree of scene evidence on a given trial scaled with the vividness with which a memory could be retrieved. This extends prior evidence showing that activation, particularly in the PhC, is stronger when scenes are recollected more vividly and in greater detail (24–28). Our data thus further validate the use of classifier evidence as a marker of memory reactivation. We expected that intentional attempts to prevent retrieval should lead to reduced scene reactivation. This was the case in the current study across the grey matter as well as for the left PhC (see also 10, 31, 32). The data thus suggest that participants successfully suppressed unwanted retrieval.

Suppression also had a sustained impact on the avoided memories: It diminished their vividness on a subsequent test. This finding adds to the extant literature by showing that suppression does not only affect the objective availability of a memory in an all-or-none fashion (4). By reducing the vividness of the memories, it also gradually diminishes the subjective quality of the information that can be retrieved. Such graded forgetting can be beneficial, for example when it allows for the continued conscious access to an aversive past event while dampening its affective impact ((2, 63), see also (62, 64, 65), for dissociations of retrieval success versus vividness).

Critically, the sustained fading of a suppressed memory has been argued to result from a deterioration of its neural representation (5, 34). This deterioration, in turn, would lower the potential of these representations to be reinstated on later retrieval attempts. To test this hypothesis, we tracked changes in the reactivation of suppressed representations.

The classifier analysis yielded evidence for a greater reduction of scene reactivation for suppressed than for baseline memories. This was the case at a global level across the grey matter as well as locally in the PhC. Moreover, consistent with the contribution of the PhC to mnemonic vividness (26), participants who displayed a greater suppression-induced reduction in scene reactivation in the PhC also experienced a greater reduction in vividness. Indeed, disruptions of cortical memory representations, such as in the PhC, may particularly lead to these graded effects of forgetting (10). That is, as cortical representations get progressively weakened, they may become increasingly susceptible to interference from overlapping representations of similar memories (66).

By comparison, hippocampal representations are encoded in a more orthogonal fashion (60, 67) and may thus be largely protected from interference. The disruption of these representations would thus not manifest as graded forgetting but eventually lead to holistic forgetting in an all-or-none fashion (23, 65, 66). Indeed, a previous study did not obtain evidence for lingering effects of suppression on hippocampal reinstatement when the memories could still be recalled (68).

A pattern classifier is a powerful tool for inferring the reactivation of category-specific neural representations (69–72). As a downside, it does not provide evidence for the reinstatement of individual exemplars from within a category. Weaker scene evidence during the retrieval of suppressed memories may thus not only reflect reduced reactivation of the respective scene. Instead, it could conceivably also result from greater reactivation of additional, non-specific scene information during the retrieval of baseline memories. However, this interpretation is difficult to reconcile with the observation that, across participants, a greater suppression-induced reduction in scene reactivation was associated with a stronger decline in vividness.

In a complementary set of analyses, we further used RSA to track the reinstatement of individual memory representations. Specifically, given that the retrieval of a memory should reinstate its representation, we expected similar activity patterns to emerge for a given memory on the pre- and on the post-test (18, 29, 73, 74). This was the case for the baseline condition, where the activity patterns were more similar for the comparison of a memory with itself than with the other memories. By contrast, the reinstatement was not significant for suppressed memories. However, we did not obtain the critical evidence that it was weaker than the baseline effect.

The absence of a significant difference in our study might simply reflect lower power for the more fine-grained, condition-rich analysis of individual activity patterns than for the more generic classifier (56). It may also result from the repetition of the identical objects as retrieval cues during the pre- and post-tests. Though the PhC is more sensitive to scenes than objects (20), the repeated objects could have contributed to the similarity of the activity patterns. This would have been the case in both the baseline and suppress condition. Such a common effect driven by the object repetitions may have obscured any condition-specific effect driven by differences in scene reinstatement.

However, the absence of an overall effect might also reflect the varying degrees to which participants were successful at suppression. Indeed, right parahippocampal reinstatement was particularly affected in those people who also experienced the strongest decline in vividness. This finding is particularly noteworthy as it corroborates the association that we had obtained with the more general index of scene reactivation. Together, the two sets of analyses thus converge in their support of our hypothesis that suppression leads to a sustained reduction in neural reinstatement.

Computational modelling suggests that suppression deteriorates memory representations via a targeted inhibition of the respective representation’s strongest, i.e., most active, features (10, 35). These simulations imply that neural representations need to be at least partially reactivated to become liable to disruption (75–77).

Indeed, during initial suppression attempts, unwanted memories often involuntarily start intruding into awareness (8, 13, 78, 79), indicating that they were partly reactivated. Such intrusions then become less frequent over time with repeated suppression attempts. The decrease in intrusions has been associated with a mechanism of reactive inhibitory control that is mediated by an upregulation of the dlPFC and a negative top-down modulation of hippocampal activation (8, 78). This inhibitory signal may be relayed via the thalamic reuniens nucleus (80) and there is some evidence that it relies on GABAergic activity in the hippocampus (81).

This account is reminiscent of the non-monotonic plasticity hypothesis which proposes that memories get weakened if they are moderately activated – irrespective of any intention to forget (35, 76, 82–85). On a neurophysiological level, this effect is reflected in long-term depression (i.e., synaptic weakening) following moderate postsynaptic depolarization (82, 86). Suppression may complement such a passive learning process with a top-down process mediated by the dlPFC. By disrupting hippocampal retrieval, the top-down process may keep the reactivation of a memory to a moderate activity level and thus render its representation amenable to synaptic weakening.

To conclude, the current study set out to examine the neural consequences of suppression that underlie the phenomenon of suppression-induced forgetting (4, 34, 36). We demonstrated that suppression rendered memories less vivid and, at the same time, provided evidence that it hindered the reactivation of their neural representations. Notably, a weaker reinstatement of the memories was also associated with a greater reduction in vividness. We thus tie the sustained phenomenological changes induced by suppression to their neural basis.

## Methods

### Participants

Thirty-seven right-handed volunteers participated in this study. They were all drawn from the participant database of the Max Planck Institute for Human Cognitive and Brain Sciences, reported no history of psychiatric or neurological disorder, gave written informed consent as approved by the local research ethics committee, and were reimbursed for their time. Four participants were excluded either due to technical problems (2), non-compliance with the instructions as assessed by a post-experimental questionnaire derived from (87) (1), or drop out (1). We thus included 33 participants in the analysis (age: *M* = 24.85 y, *SD* = 2.14 y; 17 female, 16 male). We had aimed for a final sample of 30 participants and thus recruited 37 participants in anticipation of possible exclusions due to non-compliance or excessive movement. This target sample size was chosen to exceed previous studies on suppression and based on our behavioral study.

### Materials

The stimuli for the experimental procedure were taken from (37). They comprised 60 object-scene pairs: 48 critical pairs and 12 filler pairs. The scenes were negative images depicting aversive scenes and were originally selected from the International Affective Picture System (88) and online sources. The objects were photographs of familiar, neutral objects taken from (89). Specifically, each object was chosen to resemble an object that was also part of its paired scene (but not essential to the gist of the scene). Throughout the experiment, all images were presented on a grey background. The 48 critical pairs were divided into three sets that were matched on salience of the objects, as well as the emotional valence and arousing nature of the scenes (37). Assignment of the sets to the three conditions was counterbalanced across participants.

The task for training the pattern classifier was based on (35). It included black and white photographs of five different categories: aversive scenes, neutral scenes, morphed scenes, objects, and fruits. The aversive scenes were different items taken from the same databases as the critical items. We created morphed pictures of the aversive scenes with the procedure described by (46). The morphed pictures retain the low-level visual features of the original pictures while ensuring that their content can no longer be recognized. We placed the pictures of the objects and fruits on top of phase-scrambled versions of the scenes and thus ensured that the images had the same size and rectangular shape. All images for the classifier training were normalized with respect to their luminance using the procedure described by (31). The experiment was presented using Psychtoolbox (90, 91).

### Procedures

#### Experimental design

We tested the impact of suppression on memory reinstatement using an adapted version of the Think/No-Think procedure developed by (37). This procedure entailed four phases: an initial study phase, a pre-test, the suppression phase, and a post-test. These were followed by a classifier training task in the scanner and an additional memory task (see Supplementary material). The entire session took around four hours.

During the initial study phase, participants encoded all object-scene associations. First, they saw each object-scene pair, and tried to intentionally encode the associations and, in particular, the scenes in as much detail as possible. Each pair was presented for 6 s followed by a 1 s inter-trial-interval (ITI). Following initial encoding, we presented each object as a cue and asked participants to indicate within 5 s, via button press, whether they could fully recall the associated scene. Once they had pressed the button, they had again 5 s to choose the correct scene out of an array of three different scenes (all of which were drawn from the actual stimulus set). The correct object-scene pair was then presented as feedback. This procedure was repeated up to three times until participants had correctly identified at least 60% of the scenes. To facilitate learning, this phase was split into two parts, each with half of the object-scene associations. Finally, participants were again shown all objects and asked once more to indicate whether they could recall the complete scene without feedback.

Participants then moved to the MRI scanner. Here, they saw all pairs a last time for 1.5 s each with an 800 ms ITI. The extensive learning regimen and this refresher immediately prior to the critical parts of the experiment ensured that participants had encoded strong associations and were able to vividly recall the scenes. However, it made it less likely that suppression would induce absolute forgetting rather than gradual fading of the memories (8).

During the pre-test, we presented all 48 reminders on the screen for 3 s each. Participants were asked to covertly recall the associated scene in as much detail as possible for the duration of the whole trial. They then had 3 s to rate the vividness of their recollection on a scale from 1 (not vivid at all) to 5 (very vivid). We presented no feedback at this stage. The rating was followed by a long ITI of 14 s. With this long ITI, we optimized our ability to detect the activity pattern associated with the recollection of a given scene with little contamination of the subsequent trial (35). The order of trials was pseudorandomized with at most three objects from the same condition presented in a row.

The suppression phase consisted of five blocks. During a block, each object was presented two times for 3 s. A green frame around an object indicated the *recall* task. That is, here participants were asked to recall the associated scene as vividly as possible. By contrast, a red frame around an object indicated the *suppress* task. Here, participants were asked to engage a mechanism that we have previously shown to disrupt hippocampal retrieval (7, 9, 10, 13). That is, they tried to avoid the associated scene from coming to mind while focusing on the object on the screen. If the scene were to intrude into their awareness, they had to actively push it out of their mind. Importantly, a third of the objects were not shown during this phase. These items served as baseline memories to assess weakening due to the mere passage of time. The ITIs were optimized with optseq (https://surfer.nmr.mgh.harvard.edu/optseq/) and ranged from 2 s to 8.5 s with a mean of 3 s. Participants received extensive training and feedback on this procedure on the filler memories prior to entering the scanner. Immediately following the suppression phase, participants performed the post-test. This phase was identical to the pre-test but with a different pseudorandom presentation order.

Finally, participants engaged in a classifier training task (modelled on 35) to obtain a neural pattern associated with the perception of aversive scenes. We presented pictures of the five categories in separate task blocks. During each block, they saw ten different pictures of the given category for 900 ms with a 100 ms ITI. Six of these pictures were randomly repeated within each block, thus resulting in 16 trials. Participants had to indicate the occurrence of these repetitions via a button press to ensure that they attended to the stimuli. Each category was presented in six blocks (for 30 blocks in total) in a pseudorandom presentation order with no more than two blocks of the same category in a row and with 10 s inter-block-intervals. After scanning, participants completed an additional memory task (see Supplementary material).

Participants also completed a number of questionnaires. In addition to assessing compliance with the instructions, these were designed to assess their strategy use and subjective ratings on recall and suppression success. Further, they filled in Beck’s Depression Inventory II (92), the Thought Control Ability Questionnaire (93) and the State-Trait Anxiety Inventory (94). These data were not analyzed for the current purpose.

#### fMRI data acquisition

We used a 3T Siemens Prisma MRI Scanner with a 32-channel head coil at the Max Planck Institute for Human Cognitive and Brain Sciences. Structural images were acquired with a T1-weighted MPRAGE protocol (256 sagittal slices with interleaved acquisition, field of view = 240 mm by 176mm, 1 mm isotropic voxels, TR = 2300 ms, TE = 2.98 ms, flip angle = 9°, phase encoding: anterior-posterior, parallel imaging = GRAPPA, acceleration factor = 2). Functional images were acquired using a whole brain multiband echo-planar imaging (EPI) sequence (field of view = 192 mm by 192mm, 2 mm isotropic voxels, 72 slices with interleaved acquisition (angled 15° towards coronal from AC-PC), TR = 2000 ms, TE = 25 ms, flip angle = 90°, phase encoding: anterior-posterior, MF = 3) (95, 96). 369 volumes were acquired in pre- and post-tests, 197 volumes in each suppression block and 395 volumes in the classifier training. The first five volumes of each run were discarded to allow for T1 equilibration effects. Pulse oxymeter data were collected on participants’ left hand. Participants gave their responses via a 5-button box with their right hand.

### Analyses

#### fMRI data preprocessing

The MRI data were first converted into the Brain Imaging Data Structure (BIDS) format (97). All data preprocessing was performed using the default preprocessing steps of fMRIPrep 1.5.0rc2, based on Nipype 1.2.1. (98): The respective T1 volume was corrected for intensity non-uniformity and skull-stripped, before it was segmented into cerebrospinal fluid (CSF), white matter (WM), and grey matter (GM). It was then spatially normalized to the ICBM 152 Nonlinear Asymmetrical template version 2009c using nonlinear registration.

The functional data were slice-time corrected, motion corrected, and corrected for susceptibility distortions using fMRIPrep’s fieldmap-less approach. They were then coregistered to the corresponding T1 image using boundary-based registration with six degrees of freedom. Physiological noise regressors were extracted to allow for component based noise correction. Anatomical CompCor components were calculated within the intersection of the subcortical mask and the union of CSF and WM masks, after their projection to the native space of each functional run. Framewise displacement was also calculated for each functional run. For further details of the pipeline, including the software packages used by fMRIPrep, please refer to the online documentation (https://fmriprep.org/en/20.2.0/). Our univariate analyses were performed in MNI space (following smoothing with a Gaussian kernel of 6 mm FWHM), whereas the multivariate pattern analyses (MVPA) were done on unsmoothed data in native space.

#### Regions of interest

We manually segmented the PhC on the individual T1-weighted structural images, following the anatomical demarcation protocol by (52, 53). Specifically, we defined the PhC as the posterior third of the parahippocampal gyrus (29). We further used the individual grey matter masks, segmented using FSLfast (in the fMRIPrep pipeline), as an ROI.

#### First-level fMRI analysis

Data were analyzed using SPM12 (https://fil.ion.ucl.ac.uk/spm). We decomposed the variance in the BOLD time series using general linear models (GLM) (99). For the univariate analysis of the suppression phase, we analyzed the data with a GLM including a regressor for the trials of the recall condition and a regressor for the trials of the suppress condition.

For our multivariate pattern analyses (MVPA), we assessed the individual activity patterns adopting a least-squares-single approach (100). That is, for the pre- and post-test, we estimated separate GLMs for each trial with a regressor for that specific trial and a second regressor for all other trials. For the suppression phase, a given GLM included a regressor coding for all repetitions of the same object and a second regressor for all other trials. For the classifier training task, we estimated separate GLMs for each block with a regressor for that specific block and a second regressor for all other blocks.

All of these regressors coded for the respective 3 s of each trial (or 16 s of each block for classifier training) and were convolved with the canonical hemodynamic response function. In addition, each GLM included six head motion parameters, framewise displacement, the first six aCompCor components and a block regressor as nuisance regressors. We then applied a 128-Hz high-pass filter to the data and the model. For the MVPA analyses, the resulting parameter estimates were transformed into *t*-values via a contrast of the respective individual trial versus all other trials.

#### Classification analysis

We performed the classifier analysis using the decoding toolbox (45). Specifically, we trained a linear support vector machine for each participant to distinguish activity patterns associated with intact aversive scenes versus their morphed versions. We employed a leave-one-out cross-validation approach that used, on each iteration, eleven of the twelve blocks as training data. This procedure assigns a linear weight to each voxel that reflects its importance in discriminating the two classes, thus creating a weight map. We then used the transformed weight pattern (101) to estimate reactivation as the degree of scene evidence during each trial of the pre-test, post-test, and suppression phase. This was done by calculating the dot product of the weight pattern and the respective individual *t*-map.

#### Representational Similarity Analysis

We examined the reinstatement of unique memory representations using representational similarity analysis (RSA). Specifically, we assessed whether the retrieval of a given scene was associated with a similar neural activity pattern before and after the suppression phase. This analysis used the RSA toolbox (55). It was based on the 48 trials from the pre-test and the post-test. We computed the similarity values using Pearson correlation across all voxels of the respective ROI (54). Specifically, we assessed the similarity of each item with itself (same-item similarity) and the average similarity of the item with all 15 other items from the same condition (different-item similarity) (56). By constraining the different-item similarity to items of the same category, we ensure that any differences with the same-item similarity do not simply reflect general condition differences (i.e. systematic pattern differences for baseline versus suppress items). The similarity estimates were then Fisher-transformed and averaged for each condition within subjects. We determined the magnitude of pattern reinstatement as the difference score between same-item and different-item similarity.

#### Statistical Analyses

Statistical tests were done with R version 4.0.3 (102). Repeated measures ANOVAs were conducted with the afex package (Type 3 sums of squares; 103) and effect sizes are reported as generalized eta squared. Follow-up tests were based on estimated marginal means (emmeans package, 104) using pooled variances and degrees of freedom (based on the Welch–Satterthwaite equation). The significance level was set to 5%. The robust skipped Spearman’s correlations were estimated in Matlab 2017b (105) using the robust correlation toolbox (106).

## Ethics

Human subjects: The Ethics committee of the Medical Faculty, University of Leipzig, Germany, approved the study (protocol number 333/16-ek). Participants provided written informed consent to participate in the study and for group results to be published in a scientific journal.

## Data and code availability

Because participants did not give consent for their raw functional MRI data to be released publicly within the General Data Protection Regulation 2016/679 of the EU, these data can only be made available on request to the corresponding author. Aggregated fMRI data of the Think/No-Think phase are available at https://neurovault.org/collections/KAZGAACE/. A full R Markdown of the analyses and Matlab scripts are available on OSF (https://osf.io/27dkh/?view_only=8ec25028113941f683f31703ab580533).

## Funding

This research was funded by a Max Planck Research Group awarded to RGB. The funder had no role in study design, data collection and interpretation, or the decision to submit the work for publication.

## Author contributions

A-KM and RGB designed the study. A-KM collected the data and wrote the analysis code. A-KM and RGB analyzed the data and wrote the paper.

## Acknowledgements

We thank Philipp Paulus for help in implementing the classifier training task, Roxanne Eisenbeis, Johanna Fiebig, Leonie Kanne, and Sarah-Lena Schaefer for assistance in data acquisition, data transcription and scoring, Philipp Paulus, Heidrun Schultz, Hanna Stoffregen, and Angharad Williams for comments on a draft of the manuscript, as well as Bernhard Staresina for advice on the manual PhC segmentations.

## Notes

### Competing Interest Statement

The authors have declared no competing interest.

