## Supplementary Material for "Suppression weakens unwanted memories via a sustained reduction of neural reactivation"

**Supplementary Table 1.** Control regions: *Suppress > Recall*

| Region | ~BA | Hemi-sphere | Voxels | MNI | | | t-value |
|---|---|---|---|---|---|---|---|
| | | | | x | y | z | |
| Occipital, inferior temporal | 18 | L | 10034 | -28 | -88 | 10 | 10.83 |
| | 19 | L | | -46 | -80 | -8 | 10.26 |
| | 19 | L | | -30 | -90 | 20 | 9.79 |
| Occipital, inferior temporal | 18 | R | 9037 | 40 | -90 | 4 | 9.03 |
| | 37 | R | | 36 | -44 | -18 | 8.82 |
| | 7 | R | | 30 | -68 | 30 | 8.68 |
| Superior & middle frontal * | 6 | L/R | 10089 | 8 | 22 | 62 | 8.89 |
| | 13 | R | | 28 | 20 | -12 | 8.59 |
| | 6 | R | | 48 | 6 | 28 | 7.84 |
| Thalamus | | L/R | 681 | 2 | -6 | 18 | 6.54 |
| | | R | | 4 | -22 | 18 | 5.48 |
| | | R | | 12 | -14 | 18 | 5.07 |
| Inferior frontal | 13 | L | 723 | -36 | 20 | -4 | 6.51 |
| | 13 | L | | -30 | 20 | -12 | 6.51 |
| | 45 | L | | -42 | 22 | 8 | 6.51 |
| Middle frontal | 6 | L | 738 | -48 | 4 | 30 | 6 |
| | 6 | L | | -56 | 8 | 44 | 5.12 |
| | 6 | L | | -48 | 14 | 42 | 4.95 |
| Orbitofrontal | 47 | L | 166 | -34 | 46 | -14 | 4.88 |
| | 47 | L | | -46 | 44 | -14 | 4.5 |
| Superior & middle frontal | 9 | L | 219 | -24 | 46 | 34 | 4.59 |
| | 10 | L | | -24 | 50 | 26 | 4.59 |
| | 10 | L | | -30 | 56 | 20 | 4.03 |

*Note.* Thresholded at $p < .05$, FWE cluster corrected, with a cluster forming threshold of $p < 0.001$; local maxima more than 8mm apart, minimum of 10 voxels. BA = Brodmann area. * includes peak for direct suppression by Benoit & Anderson (2012)

**Supplementary Table 2.** Modulated regions: *Suppress < Recall*

| Region | ~BA | Hemi-sphere | Voxels | MNI x | y | z | t-value |
|---|---|---|---|---|---|---|---|
| Cuneus, precuneus, HPC, amygdala, PhC | 19 | L/R | 31631 | -10 | -82 | 34 | 9.88 |
| | 23 | L | | 12 | -56 | 24 | 9.85 |
| | 18 | L | | -18 | -62 | 22 | 9.34 |
| Cerebellum | | L | 185 | 12 | -52 | -62 | 5.59 |
| | | L | | 22 | -48 | -52 | 5.22 |
| | | L | | 18 | -58 | -48 | 4.2 |

*Note.* Thresholded at $p < .05$, FWE cluster corrected, with a cluster forming threshold of $p < 0.001$; local maxima more than 8mm apart, minimum of 10 voxels. BA = Brodmann area, HPC = Hippocampus, PhC = Parahippocampal cortex.

### Retrieval practice

Repeatedly retrieving a memory facilitates subsequent recall attempts (Karpicke & Roediger 2008, Karpicke & Blunt 2011, Roediger & Butler 2011). Here, we examine the effects of such retrieval practice by comparing the *recall* and *baseline* conditions (Supplementary Fig.1).

#### *Retrieval practice preserves vividness*

Memories of the recall condition seemed to remain more vivid than those of the baseline condition from the pre- to the post-test (Supplementary Figure 1a). We tested this effect with a repeated measures ANOVA with the factors condition (recall, baseline) and time of test (pre, post). The effect of condition was significant ($F(1,32) = 5.38$, $p = .027$, $\eta^2G = .010$) and, critically, so was the interaction between condition and time of test (pre, post) ($F(1,32) = 5.53$, $p = .025$, $\eta^2G = .007$). Indeed, the conditions only exhibited a significant difference after ($t(32) = -2.90$, $p = .01$, d = -0.51) but not before ($t(32) = -0.32$, $p = .75$, d= -0.06) retrieval practice. Further follow-up tests revealed a non-significant trend for a decrease in vividness for *baseline* memories ($t(32) = 1.79$, $p = .08$, d = 0.32) with no evidence for a change for *recall* memories ($t(32) = -1.60$, $p = .12$, d = -0.28).

We obtained the same pattern in our behavioral study, with a significant effect of condition ($F(1,28) = 9.60$, $p = .004$, $\eta^2G = .028$) and a significant interaction between condition and time-of-test ($F(1,28) = 15.83$, $p < .001$, $\eta^2G = .014$). Again, conditions did not significantly differ on the pre ($t(28) = -0.9$, $p = .37$, d = -0.17) but on the post-test ($t(28) = -4.19$, $p < .001$, d = -0.79). Here, further follow-up tests indicated a decrease in vividness for baseline ($t(28) = 3.41$, $p = .002$, d = 0.64), but not for recall memories ($t(28) = -0.29$, $p = .77$, d = -0.06).

#### ***Effects on scene reactivation***

Successful retrieval is accompanied by reinstatement of the brain activity pattern that was present during encoding (Xue et al. 2010, Ritchey et al. 2013, Wing et al. 2015). However, there is mixed evidence how repeatedly retrieving a memory alters this neural representation. Some evidence indicates that it remains stable over time (Ferreira et al. 2019), whereas other studies note that it becomes more differentiated (Karlsson Wirebring et al. 2015, Hulbert & Norman 2015, Ye et al. 2020, Liu et al. 2020). Globally across the whole brain, the change in scene evidence from the pre- to the post-test did not differ for *recall* versus *baseline* memories ($F(1,32) = 2.00$, $p = .167$, $\eta^2G = .001$).

By contrast, the analysis of the PhC data, with the additional factor hemisphere (left, right), yielded a significant interaction between time of test and condition ($F(1,32) = 9.25$, $p = .005$, $\eta^2G = .003$) (and also a significant main effect of time of test ($F(1,32) = 4.83$, $p = .035$, $\eta^2G = .016$), a main effect of condition ($F(1,32) = 9.97$, $p= .003$, $\eta^2G = .006$) and an interaction between time of test and hemisphere ($F(1,32) = 4.9$, $p= .034$, $\eta^2G = .003$)) (Supplementary Figure 1b). The interaction of time of test and condition reflected a relative decrease in scene evidence for *recall* memories ($t(32) = 2.72$, $p = .01$, d = 0.48 ), with no evidence for a difference for *baseline* memories ($t(32) = 1.38$, $p = .18$, d = 0.24 ).

#### ***Relationship between retrieval-practice effects on vividness and PhC scene reactivation***

We further examined the relationship between the effects of retrieval practice on vividness and on scene reactivation in the PhC. We therefore derived, for each measure, an index of retrieval practice by subtracting the temporal difference score (pre - post) of the *baseline* condition from the difference score of the *recall* condition. On both of these indices, a more negative value thus indicates a greater practice effect. The two indices were indeed positively correlated for the left PhC as indicated by a skipped Spearman's correlation of $r = .34$, 95%-CI = [.01 .59] (Supplementary Figure 1c). People who showed a greater retrieval-induced increase in vividness yielded less of a reduction – or even an increase – in scene evidence. However, this effect was not present in the right PhC (skipped Spearman's correlation: $r = -0.07$, 95%-CI = [-.46 .35]).

#### ***Effects on reinstatement of individual representations***

A rANOVA with the factors identity (same, different), condition (recall, baseline) and hemisphere (left, right), yielded evidence for overall significant pattern reinstatement ($F(1,32) = 16.49$, $p < .001$, $\eta^2G = .006$). This effect did not interact with condition ($F(1,32) = .03$, $p= .862$ , $\eta^2G < .001$). Notably, similar to the classifier results, a greater retrieval-practice effect on vividness was accompanied by a greater practice effect on reinstatement in the left PhC (i.e., reinstatement$_{recall}$ – reinstatement$_{baseline}$) (skipped Spearman's correlation: $r$ = .43, 95%-CI = [.10 .69]) but not in the right PhC (skipped Spearman's correlation: $r$ = .28, 95%-CI = [-.09 .59]) (Supplementary Figure 1d).

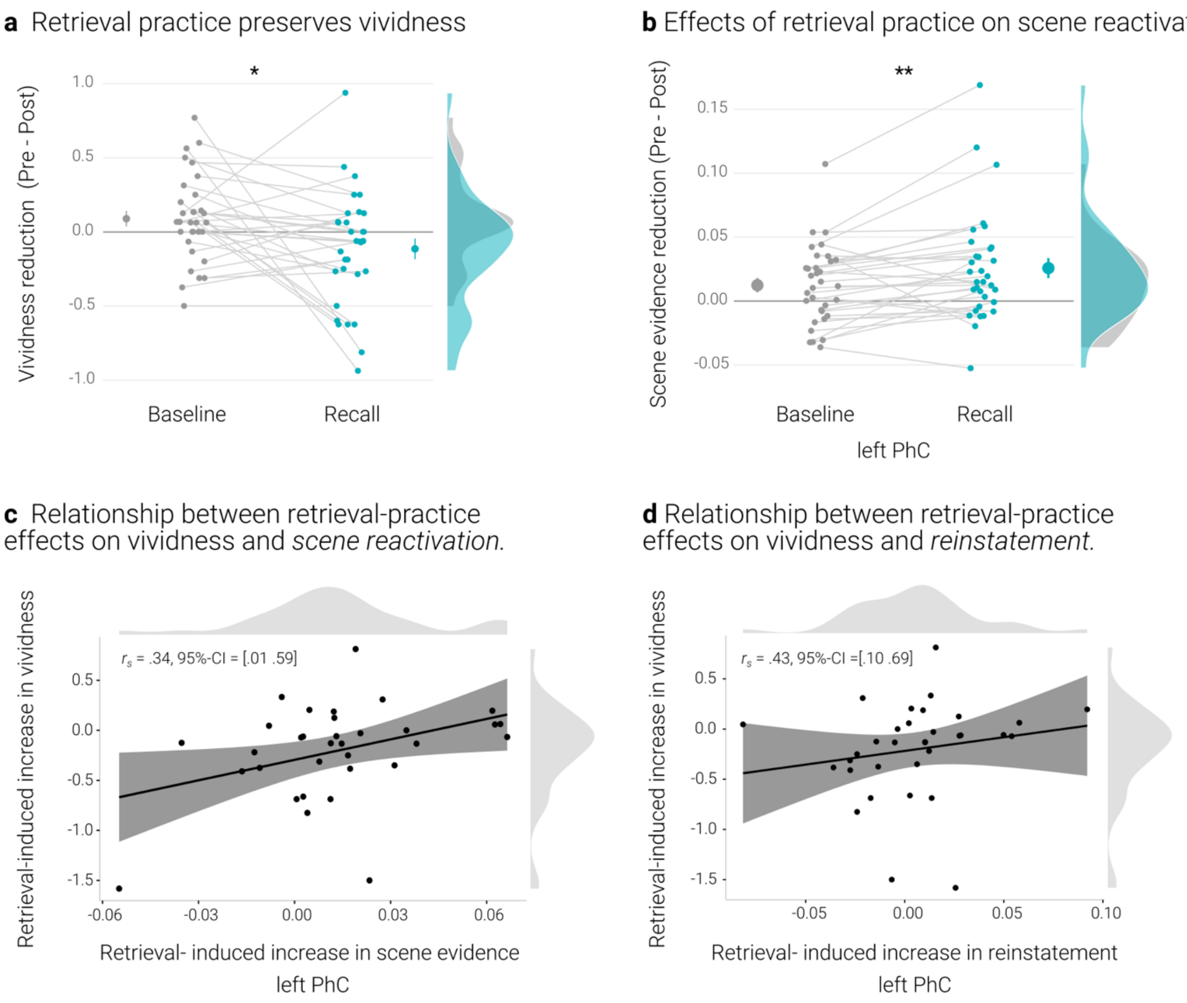


**Supplementary Figure 1. Effects of retrieval practice. a)** Retrieval retained vividness from the pre- to post-test as compared with the baseline condition. **b)** Retrieval induced a reduction in scene evidence in left PhC: scene evidence decreased from the pre- to the post- test for *recalled* memories but not for *baseline* memories. **c)** A greater above-baseline increase in vividness is associated with a greater above-baseline increase in scene evidence in left PhC as indicated by a robust skipped Spearman's correlation**. d)** This greater above-baseline increase in vividness is also associated with a greater above-baseline increase in reinstatement in left PhC**.** Larger dots indicate the mean, error bars the standard error of the mean. Black lines indicate linear regression lines, dark grey shades indicate 95% - confidence intervals, PhC: parahippocampal cortex. ** $p < .01$, * $p < .05$, n = 33.

### *Summary*

The comparison of *recall* and *baseline* memories yielded the typical behavioral effect of retrieval practice (Karpicke & Roediger 2008, Karpicke & Blunt 2011, Roediger & Butler 2011). In contrast, the analyses of the neural effects were somewhat inconclusive. Only the scene evidence obtained from the PhC showed a differential change for the practiced memories: compared to *baseline* memories, they exhibited reduced evidence for general scene reactivation. This is reminiscent of previous evidence that retrieval practice induces a greater change in parietal representations of memories that are later remembered than in those that are later forgotten (Karlsson Wirebring et al. 2015, Liu et al. 2020).

At the same time, a greater behavioral retrieval-practice effect was associated with both, a lesser reduction in scene reactivation and a more consistent reinstatement of individual memory representations. These associations suggest that retrieval-practice may facilitate the retention of a memory by stabilizing its neural representation (Ferreira et al. 2019).

Overall, the obtained pattern may indicate that retrieval practice can boost the retention of a memory in opposite fashion: on the one hand, by stabilizing its neural representation with high fidelity and, on the other hand, by changing the representation as a consequence of an ongoing semanticization (Ferreira et al. 2019) or in an attempt to reduce interference from competing memories (Kuhl et al. 2011, Hulbert & Norman 2015).

### Suppression-induced forgetting

This study set out to examine whether suppression affects the reactivation of avoided memory traces in a sustained fashion. We therefore adapted the *Think/No-Think* procedure by Kuepper et al (2014) with the insertion of additional retrieval tests just before and after the suppression phase. This procedure further strengthens the memories (Karpicke & Roediger 2008, Karpicke & Blunt 2011, Roediger & Butler 2011) compared with the original procedure and thus made it less likely that suppression would induce absolute forgetting rather than a gradual decline in the vividness of the memories (Benoit et al. 2015, Ritvo et al.  2019).

Nonetheless, as in the original procedure, participants in our study completed a final recall test. Here, they were presented with all of the objects and verbally described the associated scenes as detailed as possible within 15 s. The transcribed reports were then scored on identification, details, and gist (see Kuepper et al. 2014). On all of these measures, recall performance was numerically worse for *suppressed* than for *baseline* memories (Supplementary Table 3). However, as expected, with this modified procedure, suppression-induced forgetting (SIF) was not significant in the fMRI study (Gist: $t(32) = .33$, $p = .37$, d= 0.06 Details: $t(32)= .65$, $p = .26$, d= 0.11 Identification: W = 85, $p = .19$, d= 0.17). We obtained a similar pattern in the behavioral study with a significant effect for the identification measure only (Gist: $t(29) = 1.20$, $p = .12$, d= 0.22 Details: $t(29)= 0.88$, $p = .19$, d= 0.16 Identification: W = 70.5, $p = .038$, d = 0.35).

However, we further scrutinized the evidence for SIF across the behavioral and fMRI study. We therefore computed a mini meta-analysis in R 4.0.3 (R Core Team, 2019) using the *metafor* 2.4-0 package (Viechtbauer, 2010). Specifically, we ran a three-level random effects model, given that the three outcome measures are not independent and nested within studies (Cheung, 2014). This model accounts for the variance in the observed effect sizes (level 1), variance between effect sizes within a study (level 2), and variance between studies (level 3).

Individual effect sizes for the comparison of *baseline* and *suppress* conditions were entered as standard mean change using raw score standardization. The model parameters were estimated using restricted maximum likelihood estimation (Cheung, 2014; Viechtbauer, 2010) with the Knapp and Hartung (2003) method for calculating regression coefficients and confidence intervals. Despite our modified procedure, this analysis did yield an, albeit non-significant, trend for a small SIF effect of 0.1, $SE = .04$, $t = 2.49$, $p = .055$, 95%CI = [-0.003 0.203], with no significant heterogeneity between the studies $Q = 1.34$, $p = .93$.

**Supplementary Table 3.** Behavioral results of the final memory test

| Study | Sample size | Age in years M(SD) | Measure | Baseline M (SD) | Suppress M (SD) | Recall M (SD) |
|---|---|---|---|---|---|---|
| Behavioral Study | 30 (15 female) | 23.83 (1.76) | Identification | 0.94 (0.18) | 0.92 (0.19) | 0.94 (0.18) |
| | | | Gist | 0.52 (0.20) | 0.48 (0.19) | 0.5 (0.19) |
| | | | Details | 9.24 (2.71) | 9.07 (2.73) | 9.47 (2.84) |
| fMRI study | 33 (17 female) | 24.85 (2.14) | Identification | 0.98 (0.05) | 0.96 (0.07) | 0.96 (0.06) |
| | | | Gist | 0.55 (0.15) | 0.54 (0.14) | 0.55 (0.15) |
| | | | Details | 9.77 (1.61) | 9.66 (1.74) | 9.71 (1.75) |
